# Nested mobile genetic elements mediating antimicrobial resistance genes mobility within and between cells in clinical and environmental *Acinetobacter* isolates

**DOI:** 10.1101/2025.06.10.658816

**Authors:** Kenneth Bongulto, Ngure Kagia, Hisamichi Tauchi, Satoru Suzuki, Kozo Watanabe

## Abstract

Mobile genetic elements (MGEs) play a central role in the acquisition and dissemination of antibiotic resistance genes (ARGs). This study analyzed the distribution of MGEs using the whole-genome sequences of 38 *Acinetobacter* isolates from patient, environmental, and pig waste samples. Insertion sequence (IS) elements were identified as the most common MGE type, followed by plasmids and prophages, with significant variations depending on isolation sources. Pig waste isolates exhibited the highest mean number of plasmids, while prophages were more prevalent in environment-associated isolates. Interestingly, we observed a significant positive correlation between the number of plasmids and number of defense systems. In contrast, a significant negative correlation was identified between the number of prophages and number of defense systems. Plasmids were the primary vectors of ARGs. Co-localization of multiple ARGs, such as *msr(E)* and *mph(E)*; *bla*_OXA-58_, *tet*(*Y*), *mph*(*B*), and *sul2* within a single plasmid was observed, suggesting potential for co-resistance and multidrug resistance. ARGs were enriched in p*dif* modules, with up to 10 distinct ARGs observed within a single module. Moreover, IS elements were more densely concentrated in conjugative elements. The study further highlights the role of nested MGEs in enabling hierarchical ARG transfer processes both within and between bacterial cells. Additionally, putative novel genomic resistance islands (GRIs) were identified in non-*baumannii Acinetobacter* species, representing the first documentation of GRIs outside the *Acinetobacter calcoaceticus-baumannii* (Acb) complex. This study provides new insights into the mechanisms of ARG dissemination in *Acinetobacter*, particularly the role of MGEs in facilitating hierarchical gene transfer processes.

**IMPORTANCE:** Understanding the mechanisms behind antimicrobial resistance gene (ARG) dissemination in various environments is crucial for combating the growing global problem of antimicrobial resistance. This study investigated the distribution patterns of mobile genetic elements (MGEs) across distinct ecological contexts, analyzing samples from patients, pig wastewater, and environmental surface water. By investigating the genomes of *Acinetobacter* species from various isolation sources, we revealed that pig waste-associated isolates possessed more plasmid-borne ARGs compared to patient- and environment-associated isolates. Further, we demonstrated that ARG-bearing plasmids possessed higher density of IS elements than ARG-free plasmids. Putative novel genomic islands were also observed in non-*baumannii* species isolated from pig wastewater. These findings elucidate the hierarchical structure of ARG transfer mechanisms mediated by nested MGEs, providing valuable insights into the emergence of resistance across both clinical and environmental ecosystems.

## 1. INTRODUCTION

Horizontal gene transfer (HGT) is a fundamental driver of bacterial evolution, enabling the rapid acquisition of beneficial genetic traits that enhance fitness and survival (1). Through various mechanisms including transformation, conjugation, transduction, and vesiduction, bacteria can acquire diverse functional genes (2). Mobile genetic elements (MGEs) serve as key mediators of these processes, facilitating gene transfer both within and between cells (3, 4). These gene transfers are significant for bacterial adaptation, particularly in the development of antimicrobial resistance (5). The prevalence of MGEs, including intracellular types (e.g. insertion sequences (IS) elements, transposons, and integrons) and intercellular types (e.g. plasmids, prophages, integrative conjugative elements (ICEs), integrative mobilizable elements (IMEs), underscores their crucial role in bacterial evolution and adaptation to changing environments (6, 7). *Acinetobacter* species are well known opportunistic pathogens, inhabiting various environments such as hospital settings, human and animal wastewater and environmental waters. Infection caused by multidrug-resistant *Acinetobacter* has been a significant concern. Research focusing on the frequency of MGEs and their associated antibiotic resistance genes (ARGs) is essential to mitigate the risk of multidrug resistance development in the genus *Acinetobacter*.

While MGEs are present across all bacterial species, the frequency of different MGE types can vary significantly depending on the host bacterium’s ecology and environment. For instance, prophages, an intercellular MGE type, display variable abundance between *Staphylococcus aureus* isolates, being present in human isolates but absent in pig isolates (8). Similarly, plasmids, which are extrachromosomal MGEs, are frequently detected in livestock wastes (9, 10). The abundance of MGE types is also influenced by the presence of defense systems that control HGT (11, 12). Bacteria utilize sophisticated defense systems, such as CRISPR-Cas (13, 14) and restriction modification systems (15), which recognize specific DNA sequences to target phages and plasmids.

The prevalence of ARGs varies across different MGE types. Previous studies have shown that ARG prevalence is primarily elevated in intracellular MGEs types, such as TEs (16, 17). Among intracellular MGE types, IS elements have been shown to have higher ARG prevalence compared to integrons (7). Among intercellular MGE types, higher ARG prevalence has been reported in plasmids compared to prophages (18, 19). Plasmids can be categorized as conjugative, mobilizable, or nonmobilizable depending on the presence of conjugative genes such as relaxase and conjugative coupling proteins (20). Although plasmids are well-known ARG vectors, over 70% of plasmids lack ARGs (21). Based on these observations, we hypothesize that plasmids harbor a high frequency of ARGs due to the presence of IS elements that accumulate ARGs.

Nested MGE structures, or intracellular MGEs embedded within an intercellular MGE, mediate both intracellular and intercellular ARG transfer, forming sophisticated networks of gene transfer (22, 23, 24). One example of nested MGE structures is plasmids harboring IS elements. Since IS elements are only capable of intracellular movement, a nested MGE structure is vital for their efficient intercellular transfer. For example, IS elements can mediate the intracellular transfer of ARGs from chromosomes to plasmids, while the plasmids that receive these ARGs can mediate the intercellular transfer of ARGs. Notably, IS elements demonstrate distinct distribution patterns between genomic compartments, with plasmids showing significantly higher IS density compared to chromosomal regions (25). However, the frequency of IS elements among different types of intercellular MGEs has yet to be fully characterized.

Genomic resistance island (GRI) are variable regions in the bacterial genome formed through intracellular and intercellular ARG transfer (26). These regions integrate MGEs, including integrons and transposons, along with associated ARGs. In *Acinetobacter baumannii*, GRIs have been extensively characterized, contributing to the development of multidrug resistance by harboring different ARGs (26–34). However, while studies have documented GRIs in *A. baumannii* from several parts of the world, knowledge gaps persist regarding the types of GRIs in non-*baumannii* species. Recent investigations have identified AbaR-type GRIs in non-*baumannii* species, such as *A. nosocomialis* and *A. seifertii*, demonstrating interspecies transfer capabilities (35). Notably, a significant knowledge gap exists regarding the characteristics and prevalence of GRIs in non-*baumannii* species, particularly the pathogenic *Acinetobacter calcoaceticus – baumannii* (Acb) complex group.

The present study examines the mechanisms of ARG mobility in *Acinetobacter* species from various sources (i.e., patient, environmental, and pig waste samples), with particular emphasis on MGEs and their role in ARG acquisition and dissemination. The aim of this study is to characterize the hierarchical process of ARG transfer mediated by intracellular and intercellular MGEs. Specifically, we investigated whether the frequencies of MGE types in *Acinetobacter* species are influenced by source origins and their defense systems. Additionally, we examined the ARG prevalence among different types of MGEs and the co-localization patterns of ARGs within MGEs. Subsequently, we also analyzed the nested MGE structures and investigated which type of intercellular MGEs exhibits a high IS element frequency. Based on these results, we tested the hypothesis that ARGs are frequently present in plasmids because IS elements frequently harbor ARGs and are highly abundant in plasmids. Lastly, we discuss the intracellular and intercellular movement of ARGs and how it contributes to the formation of genomic islands in *Acinetobacter*.

## 2. MATERIALS AND METHODS

### 2.1. Data Collection

The datasets, namely plasmid sequences and chromosome sequences of *Acinetobacter* species from the different co-selection patterns of ARGs and virulence genes (36), were retrieved from the NCBI database under the BioProject accession number PRJNA1184881. All *Acinetobacter* genomes were sequenced using both Illumina and Oxford Nanopore Technology. The hybrid assembled genome of *Acinetobacter* isolates were used to extract MGE sequences. Our set of 38 isolates originated from patients, pig wastewater, municipal wastewater and natural surface water. The plasmid dataset (*n*=193) only includes circularized DNA and complete sequences. Other intercellular MGEs (e.g. prophages, ICEs, IMEs) were extracted from the chromosome sequences. Further, chromosome and plasmid sequences were used to identify intracellular MGEs, such as IS elements, transposons, and integrons.

### 2.2. Bioinformatics analysis

Plasmid mobility was predicted using the parameter mob_typer in Mobsuite v3.1.8 (37) and plasmid typing was conducted using blastn against the *Acinetobacter* Plasmid Typing (APT) scheme (38). Prophage regions in both chromosomes and plasmids were predicted using the default parameter of PHASTEST (39). Mobile Genetic Element Finder (40) was used to identify IS elements. Transposons were detected using BacAnt (41) with the minimum identity threshold of 90% and 60% coverage. Integrons were detected using the parameter –local-max in Integron Finder v.2.0 (42) to allow for a more sensitive search and minimize the false positive rate. Integrative conjugative elements and integrative mobilizable elements were predicted using the default parameter of ICEberg 2.0 (43). Genomic islands and p*dif* modules were predicted using the default parameter of IslandViewer 4 (44) and pdifFinder (45), respectively. To predict ARGs in MGE sequences, Abricate (46) was used with the CARD v3.2.4 (47) and ResFinder v.4.6.0 (48) databases with a minimum identity threshold and minimum coverage threshold of 90%. Lastly, defense genes were predicted using the DefenseFinder tool (49).

### 2.3. Gene synteny analysis

Each MGE sequence was annotated using Bakta (50) and the full database version (v.5.0) was used to obtain the best annotation results. The annotated .gbk files were used for the gene synteny analysis using the R package geneviewer v0.1.6 (51). The sequences of ARG-bearing plasmids and genomic islands in chromosome were used for the ARG synteny analysis.

### 2.4. Visualization and Statistical analysis

The frequency of IS elements, plasmids, prophage regions, integrative conjugative elements (ICEs), and integrative mobilizable elements (IMEs), as well as defense systems among patient-, pig waste-, and environment-associated *Acinetobacter* isolates were determined using Kruskal-Wallis test with post hoc Wilcoxon rank-sum test. Statistical significance was considered at *p* <0.05. Additionally, the correlation between the number of plasmids and number of defense systems, and between number of prophages and number of defense systems were examined. The number of defense systems, plasmids, and prophages were normalized using the total number of protein-coding sequences (CDS). The correlations were evaluated using the cor function in R v.4.2.0. with statistical significance set at *p* < 0.05. The significant ARG co-localization on a single plasmid was tested using Fisher’s exact test with fisher.test function in R. The test was performed for all plasmid-bearing ARGs. The upset plot for the co-localization patterns of ARGs were generated in R v.4.2.0. Kruskal-Wallis test was conducted in comparing the frequency of insertion sequence (IS) elements across different intercellular MGEs such as plasmids, prophages, ICEs, and integrative mobilizable elements (IMEs) using the function kruskal.test in R v.4.2.0. Likewise, Wilcoxon rank test was also performed to determine the significant difference in terms of IS element carriage between plasmid-bearing ARGs and plasmid without ARGs. Further, the relationship between p*dif* modules and ARG carriage in plasmid-bearing ARGs and plasmids without ARGs were investigated. Wilcoxon test was also performed with statistical significance set at *p* <0.05. Sankey plot was built using the networkD3 package in R to show the association of ARGs, intracellular MGEs, and intercellular MGEs.

## 3. RESULTS

### 3.1. Prevalence of mobile genetic elements in *Acinetobacter* isolates

Of the 38 Acinetobacter isolates examined, IS elements (*n* = 1045) were the most prevalent MGEs, followed by plasmids (*n* = 193) and prophages (*n* = 103) (see **Supplementary Fig. 1.A)**. Integrative conjugative elements (ICEs) and integrative mobilizable elements (IMEs) were the least prevalent (*n* = 28 and *n* = 13, respectively). Significant differences in the frequencies of each MGE type were observed between patient- and pig waste-associated isolates, specifically for IS elements, prophage, and plasmid carriage **(Fig. 1.A)**. However, there was no significant difference in the frequency of ICE and IME carriage among Acinetobacter isolates. IS elements were significantly more abundant in environment-associated isolates than pig waste-associated isolates (*p* = 0.0114). In contrast, plasmids were significantly more prevalent in pig waste-associated isolates than in patient- and environment-associated isolates (*p* < 0.0001). Environment-associated isolates had more prophages than pig waste-associated isolates. We also observed a significant difference in the mean frequencies of defense systems among *Acinetobacter* isolates from different sources. Pig waste-associated isolates had a higher frequency of defense systems than both environment-(*p* < 0.05) and patient-associated isolates (*p*<0.01) **(Fig. 1.B)**. Furthermore, we observed a significant positive correlation (*p*=0.0012) between the number of plasmids and the number of defense systems **(Fig. 1.C)**. Interestingly, we also found a significant inverse correlation (*p =* 0.0017) between the number of defense systems and the number of prophages in the *Acinetobacter* genomes **(Fig. 1.D)**.

**Figure 1.**
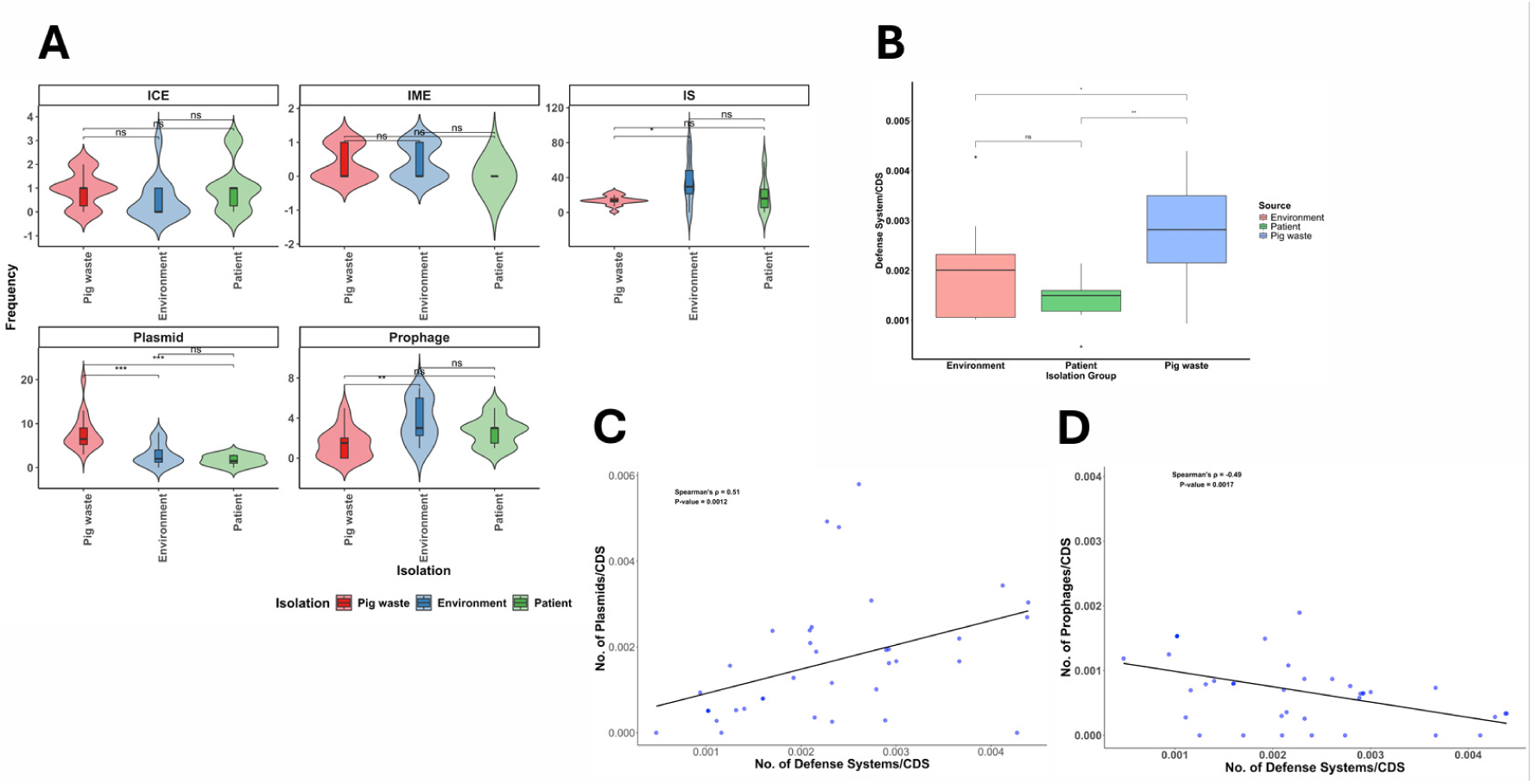
Frequency of mobile genetic elements (MGEs) and defense systems across different isolation sources. **A.** The frequency of integrative conjugative elements (ICEs) and integrative mobilizable elements (IMEs) in *Acinetobacter* chromosomes. Frequency of insertion sequence (IS) elements in the chromosome of *Acinetobacter* species (\**p* <0.05). Plasmid counts among patient-, pig waste- and environment-associated isolates (\*\*\**p* <0.001). Number of prophages integrated in the chromosome of *Acinetobacter* isolates (\*\**p* <0.01). **B.** The frequency of defense systems in *Acinetobacter* species across different isolation sources (*p\**<0.05, \*\**p*<0.01). Kruskal-Wallis test was performed to compare the average number of MGEs and defense systems among *Acinetobacter* isolates. **C.** Correlation plot of the number of plasmids and number of defense systems using Spearman’s rank correlation (R=0.51, *p*=0.0012). **D.** Correlation plot of the number of prophages and number of defense systems using Spearman’s rank correlation (R=-0.49, *p*=0.0017). Each dot represents a genome (n=38).

### 3.2. ARG distribution, co-occurrence, and enrichment in plasmids

The extent of ARG carriage in different MGE types was also investigated. Most ARGs were found to be associated with 42 plasmids and one integrative mobilizable element (IME), with no ARGs detected in prophages or ICE regions. A total of 193 plasmids were identified in the 38 *Acinetobacter* isolates. Most of the isolates’ plasmid repertoire consists of nonmobilizable plasmids (n = 105, 54.4%), mobilizable plasmids (n = 81, 42%), and conjugative plasmids (n = 7, 3.6%) (see **Supplementary Fig. 2.A)**. One hundred thirty-nine plasmids (72%) could not be assigned to a plasmid typing scheme using the rep gene (see **Supplementary Fig. 2.B)**. Similarly, relaxase were not detected in 105 plasmids (54%) using the mob typing scheme (see **Supplementary Fig. 2.C)**. Plasmid sizes exhibited remarkable diversity, ranging from 1.4 kb to 657 kb (see **Supplementary Fig. 2.D)**.

A total of 42 plasmids harbored ARGs, and the majority (n=40, 95.2%) were found in pig waste-associated isolates. Large size plasmids (>100 kb) tend to harbor more ARGs (4-16 ARGs) and were categorized as mobilizable or nonmobilizable **(Fig. 2.A)**. Of the ARG-carrying plasmids, 62 % (n = 26) were conjugative or mobilizable, while 38% (n=16) were non-mobilizable. Macrolide resistance genes (*mph(E)* and *msr(E*)) and aminoglycoside resistance genes (*aph(3”)-Ib* and *aph*(*6*)*-Id*) tended to co-occur in 39% and 32% of ARG-carrying plasmids, respectively **(Fig. 2.B)**. Further analysis revealed that 19 pairs of ARGs were co-localized on a single plasmid, showing significant associations (*p* = <0.05, using Fisher’s test) **(Supplementary Table 1).** Up to 14 different ARGs were detected on a single plasmid. Nine prophage regions were also detected in eight plasmids. However, no ARGs were directly encoded within the prophage regions of these plasmids.

**Figure 2.**
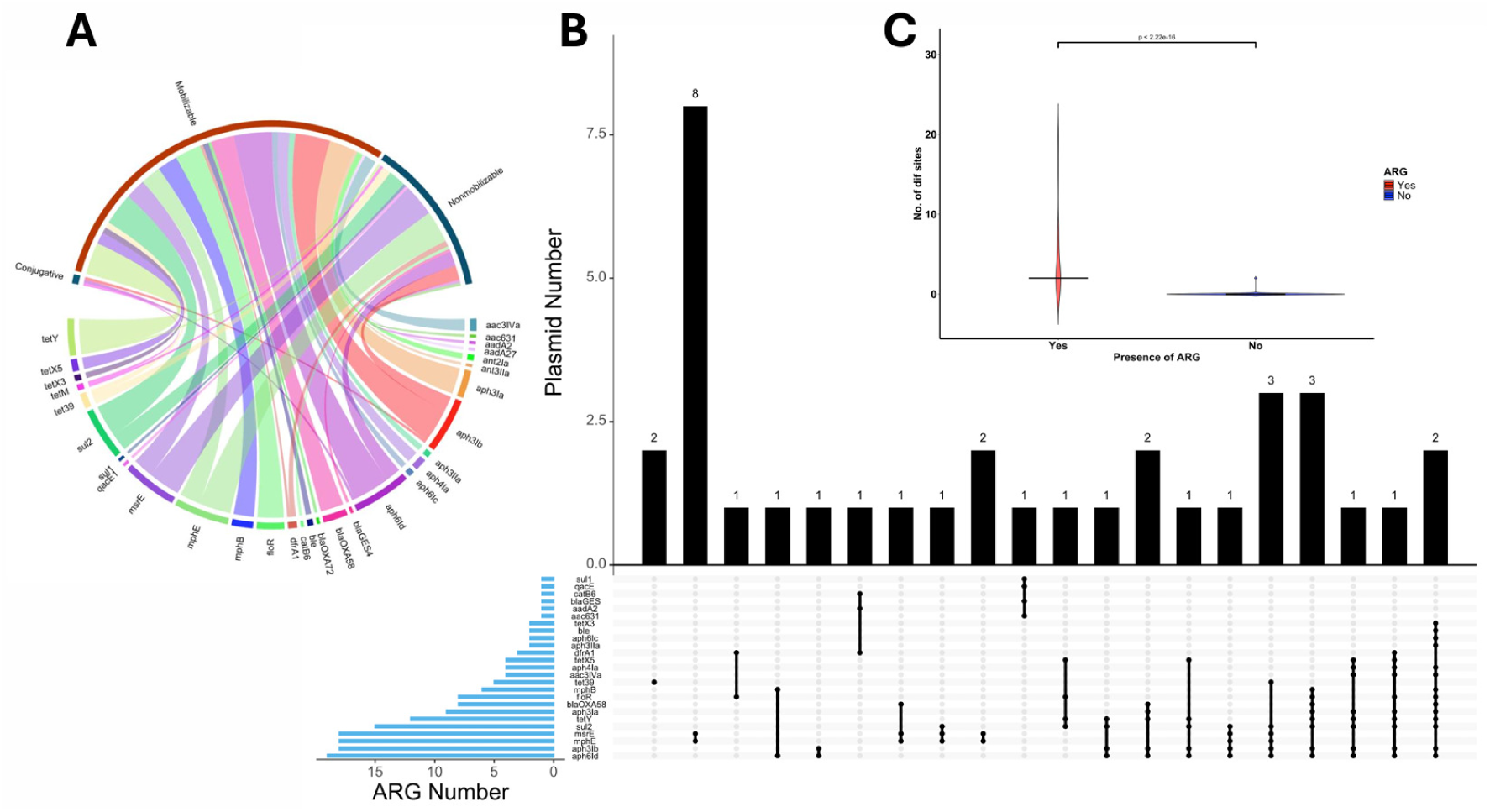
ARG distribution, co-occurrence, and enrichment in *Acinetobacter* plasmid. **A.** Chord diagram showing the association of ARGs with different plasmid types. **B.** Upset plot showing the ARG number in blue bar plots, ARG co-localization patterns shown in interconnected dots and number of plasmids having the ARG co-localization pattern in black bar plots. **C.** Comparison of the number of p*dif* sites between ARG-bearing plasmids and plasmids without ARGs (*p* <2.22e-16). Wilcoxon test was performed to compare the average number of IS elements and number of p*dif* sites between ARG-bearing plasmids (red) and plasmids without ARGs (blue).

We assessed the presence of p*dif* sites in all ARG-carrying plasmids, as well as investigating whether these sites were associated with the acquisition of ARGs. As many as 20 p*dif* sites were detected in a single ARG-bearing plasmid. Multiple ARG types were flanked by these p*dif* sites, forming ARG p*dif* modules (see **Supplementary Fig. 3.A-3.G)**. In total, seven different ARG p*dif* modules were identified: p*dif-*(*msr(E)-mph(E)*), p*dif*-(*tet*(*39*)), p*dif*-(*bla*_OXA-58_), p*dif*-(*aph(3”)-Ib – aph*(*6*)*-Id – tet(Y) – aph(3’’)-Ia*), p*dif*-(*aph(3”)-Ia – tet(Y) – aph*(*6*)*-Id – aph(3”)-Ib – floR – mph(B) – sul2*), p*dif*-(*sul2 – aph(3”)-Ib – aph*(*6*)*-Id*), and p*dif*-(*aph*(*4*)*-Ia – aac*(*3*)*-IVa – aph(3”)-Ia – aph(3”)-Ib – aph*(*6*)*-Id – tet(Y) – tet(X5) – floR – dfrA1 – sul2*). The number of ARGs in p*dif* modules ranged from 1 to 10 per module. Plasmids with ARGs had more p*dif* sites than those without ARG (*p* = 2.22 x 10^-16^, Wilcoxon test) **(Fig. 2.C)**. Furthermore, there was a significant association between the number of p*dif* sites and the number of ARGs in ARG-carrying plasmids (*p* = 2.32 x 10^-25^, Fisher’s test).

### 3.3. Nested MGE structures in *Acinetobacter*

A total of 29 types of IS elements were detected among 193 plasmids **(Supplementary Fig. 4.A)**. The most prevalent IS elements were ISAba14 (n=23) and IS17 (n=23), followed by ISOur1 (n=20), ISAba34 (n=19), and IS1006 (n=18). We examined whether ARG enrichment in plasmids is associated with the presence of IS elements. Plasmid-bearing ARGs possessed more IS elements compared to plasmids without ARGs (*p =* 0.0012) **(Fig. 3.A)**. The number of IS elements was positively correlated with the plasmid size **(Supplementary Fig. 4.B)**. The Sankey plot analysis showed that nine ARGs existed in TEs on plasmids (**Fig. 3.B)**. The aminoglycoside resistance genes, *aph*(*6*)*-Id* and *aph(3”)-Ib,* both occurred on plasmid Tn6205, while *aph(3”)-Ia* was found on Tn4352. Furthermore, ARGs such as *dfrA1*, *bla*_GES-4_, *aac(6’)-31*, *aadA2*, and *catB6* were identified within gene cassettes associated with integrons.

**Figure 3.**
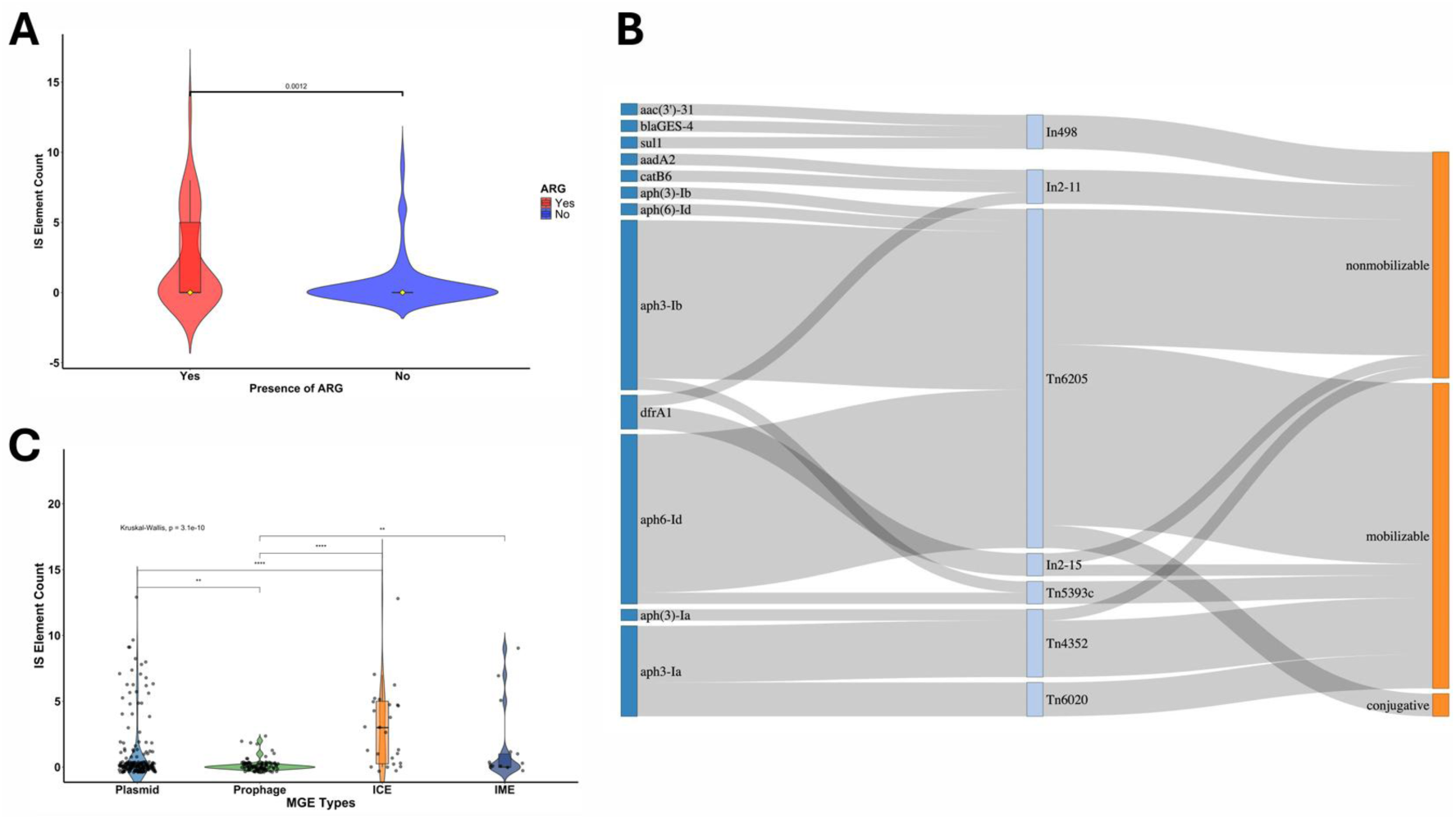
Nested MGE structures in *Acinetobacter*. **A.** Comparison of IS element carriage between ARG-bearing plasmids and plasmids without ARGs (*p* = 0.0012). **B.** Sankey plot showing the association of antimicrobial resistance genes (green) in nested MGEs - transposable elements (orange) and different plasmid types (blue). **C.** Frequency of IS elements among different intercellular MGE types (\*\**p* <0.01, \*\*\*\**p* <0.0001). Kruskal-Wallis test was performed to compare the frequency of IS elements in A and C.

Among the four types of intercellular MGEs, IS elements were predominantly found in conjugative elements, plasmids and ICEs **(Fig. 3.C)**. The frequency of IS carriage was significantly different between plasmids and prophages (*p* <0.0001), as well as between ICE and prophages (*p* <0.001). The frequency of IS elements also varies among plasmid types, with mobilizable and non-mobilizable plasmids possessing more IS elements (see **Supplementary Fig. 4.C**). Furthermore, no significant difference was observed in the frequency of IS element carriage among plasmids from different isolation sources (patient, pig waste, environment) (*p* = 0.61, Kruskal-Wallis) (see **Supplementary Fig. 4.D)**. However, the types of IS elements varied among plasmids isolated from different isolation sources, with some types being exclusively found in plasmids from a specific isolation source (see **Supplementary Fig. 5)**.

Duplication of ARGs on plasmids was also observed. Some of the ARGs that duplicated were aminoglycoside resistance genes, such as *aph(3”)-Ib, aph*(*6*)*-Id,* and *ant(2”)-Ia*. Additionally, we revealed that several plasmids from different bacterial hosts tend to have similar set of ARGs (see **Supplementary Fig. 6.A & 6.B**). We demonstrated the movement of macrolide resistance genes (*msr(E)* and *mph(E*)) and aminoglycoside resistance genes (*aph(3”)-Ib* and *aph*(*6*)*-Id*) between different plasmids in different hosts. A similar set of ARGs was also observed between co-residing plasmids. Specifically, this was observed for the *aph(3”)-Ib* and *aph*(*6*)*-Id*, *sul2,* and *msr(E)* and *mph(E)* genes (see **Supplementary Fig. 7.A-7.C)**.

Nine prophage regions, ranging in size from 44 to 183 kilobases, were identified across eight plasmids. Individual plasmids harbor between one or two prophages. Notably, these prophages were not associated with *Acinetobacter* as their host bacterium; rather, they represented various phage types, including Escher_RCS47, Stx2_c_1717, Escher_PA28, Entero_VT2phl, Stx2_II, Staphy_Spbeta, Faecal_FP_Taranis, and Salmon_epsilon15 (see **Supplementary Table 2)**. Furthermore, we observed that prophage regions in plasmids exhibited a significantly higher frequency of IS elements than those located within the chromosomes (*p =* 7.5e-13) **(Supplementary Fig. 8.A)**. Out of the nine prophages in plasmids, six were detected as intact prophages in terms of completeness. We also found that intact prophages had a significantly higher frequency of IS elements compared to incomplete prophages (*p =* 0.034) **(Supplementary Fig. 8.B)**.

### 3.4. Genomic islands in non-*baumannii* species

We detected intercellular ARG movement through plasmid-mediated mechanism and intracellular ARG transfer between co-resident plasmids and the chromosome using ARG synteny analysis. We detected four putative novel genomic islands in non-*baumannii* pig waste-associated isolates **(Supplementary Fig. 9.A-9.D)**. The ARG synteny analysis revealed that mobilizable ARG regions in plasmids of the pig waste-associated isolates showed high sequence similarity with the ARG regions in the chromosome of the host bacteria. Furthermore, transposable elements in plasmids showed homologous regions within the chromosome. The first GRI type was detected in a plasmid of *A. towneri* (EA24) flanked by IS1006 and containing a sulfonamide resistance gene, as well as cobalt, zinc, and cadmium resistance gene *czcA* gene **(Fig. 4.A)**. The second GRI type, observed in *A. towneri* (EA13), was marked by an IS3 family-bound GRI containing resistance genes (*aph(3”)-Ib, aph*(*6*)*-Id,* and *sul2*), metal resistance genes (*copA, copB, merR, zitB, czcABD*), and a toxin/anti-toxin system **(Fig. 4.B)**. The third GRI type included an integron class 2, associated with the Tn7 transposon family, and was identified in *A. towneri* (EA14), harboring a gene cassette bearing a trimethoprim resistance gene (*dfrA1*) **(Fig. 4.C)**. The fourth GRI type, observed in *A. haemolyticus* (EA17), featured an ISAba1-bound GRI that contained aminoglycoside resistant genes (*aph(3”)-Ia*, *aph(3”)-Ib*, and *aph*(*6*)*-Id*); a tetracycline resistance gene (*tet(X3)*); and sulfonamide resistance gene (*sul2*) **(Fig. 4.D)**.

**Figure 4.**
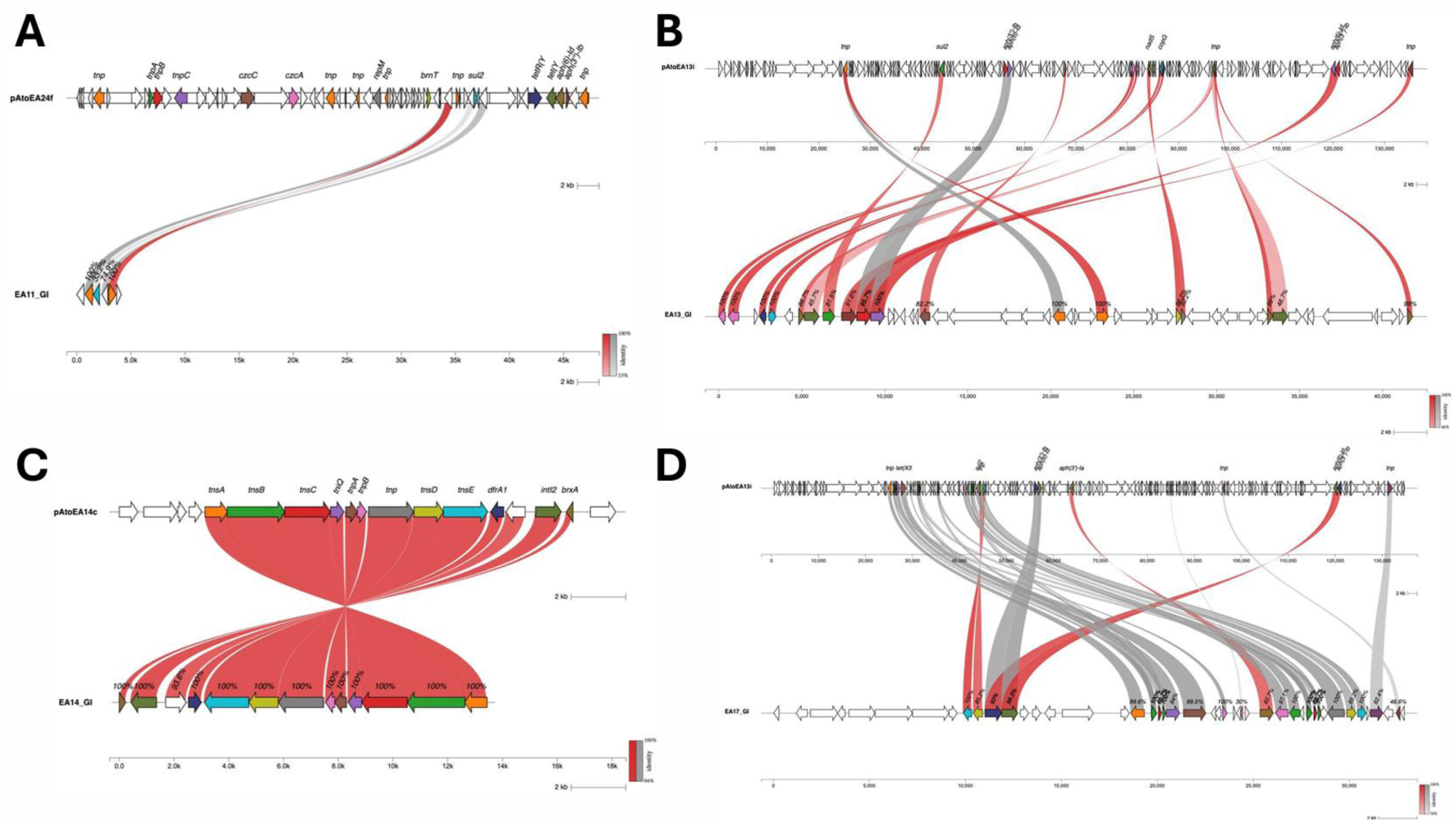
Genomic resistance islands (GRIs) in non-*baumannii* isolates. **A.** ARG synteny analysis showing homologous region of *sul2* gene between plasmid pAtoEA24f (*A. towneri*) and chromosome of *A. indicus* EA-11. **B**. ARG movement from a co-residing plasmid pAtoEA13i (*A. towneri*) into the chromosome (*A. towneri* EA-13) **C.** ARG synteny analysis showing homologous region of *dfrA1* gene in an integron of plasmid pAtoEA23i *A.* (*towneri*) and a plasmid pAtoEA14c from a different bacterial host (*A. towneri*). The same region was found integrated into the chromosome of *A. towneri* EA-14. **D.** ARG synteny analysis of ARGs from plasmid pAtoEA13i (*A. towneri*) and genomic island in *A. haemolyticus* EA-17. Red lines indicate high sequence similarity matches while gray lines indicate reverse or inverted synteny.

## 4. DISCUSSION

### 4.1. Variation in MGE frequency among *Acinetobacter* isolates depends on isolation source and defense genes

IS elements were identified as the most prevalent MGE type in the genomes of the 38 isolates analyzed, followed by plasmids and prophages. This result aligns with a previous finding from Khedkar et al. (17), which demonstrated that IS elements were the most abundant MGE type in prokaryotic genomes, such as those of Firmicutes and Proteobacteria.

A notable differentiation in the distribution of MGEs based on the isolation source was observed. Plasmids were abundant in pig waste-associated isolates, whereas they were rare in patient- or environment-associated isolates. These findings are consistent with previous studies, which have reported the widespread plasmids and demonstrated a strong positive correlation between the numbers of plasmids and ARGs per cell (52, 53). Specifically, integrons and conjugative plasmids, which are widely distributed in livestock waste, serve as key vectors for ARG dissemination via HGT (54, 55). The extensive use of antibiotics in pig farming imposes significant selective pressure, favoring bacterial populations that harbor ARG-bearing plasmids (56). These observations suggest that the high antibiotic pressure in pig waste environments promotes the selection of MGEs, particularly plasmids, which drive the dissemination of antimicrobial resistance.

In contrast, prophages were observed at a higher frequency in environment-associated isolates compared to those from patients or pig waste. This trend corresponds with prior studies that identified a high prevalence of prophages in environments such as wastewater treatment plants (WWTPs), which harbor diverse bacterial populations of human origin (57). A large-scale analysis of prophages from 13,713 complete prokaryotic genomes also identified genera such as *Acinetobacter*, *Enterobacter*, and *Pseudomonas* as major prophage reservoirs (58). Furthermore, it has been reported that prophages were frequently detected in human-derived *Staphylococcus aureus* isolates compared to those from pig origin (8).

To further investigate the factors influencing the frequency of MGEs, we examined the presence of bacterial defense systems within each *Acinetobacter* genome. Notably, a negative correlation was observed between the number of defense systems and the number of prophages, highlighting the role of defense systems in restricting prophage acquisition. This finding is consistent with findings from Kohgay et al. (59), who reported fewer phage-related genes in bacterial genomes with a higher abundance of defense systems. Similarly, Costa et al. (60) demonstrated that an increase in defense systems enhances bacterial resistance to phage infections, further supporting their involvement in limiting prophage acquisition.

Interestingly, isolates derived from pig waste exhibited the highest frequency of defense systems. Additionally, we observed a significant positive correlation between the numbers of defense systems and plasmids. Plasmids, while known as principal vectors of ARG dissemination, also carry diverse defense systems, which protect host bacteria from phage infections. Recent studies (61, 62) have emphasized the strong association between plasmids and bacterial defense systems, with plasmids playing a pivotal role in the accumulation and dissemination of these systems across bacterial genomes.

Based on these observations, we propose a hypothetical model wherein isolates originating from pig waste environments acquire plasmids carrying multiple ARGs under the selective pressure of high antibiotic use. These plasmids may additionally harbor defense systems that protect the host bacteria against phages. Future experimental studies are needed to validate this hypothesis and further elucidate the interplay between plasmids, defense systems, and prophage dynamics in different environmental contexts.

### 4.2. The extent of ARG carriage in *Acinetobacter* was predominantly found in plasmids

Our analysis revealed that ARGs in *Acinetobacter* isolates were predominantly associated with plasmids. Notably, the ARG carriage accounted for only 21% of the total plasmids identified across all isolates, consistent with former findings indicating that more than 70% of plasmids lack known resistance genes (21). Interestingly, most ARGs in this study were found in mobilizable and non-mobilizable plasmids, in contrast to other studies showing an enrichment of ARGs in conjugative plasmids (63).

Distinct patterns of ARG co-localization were observed in plasmids. For example, macrolide resistance genes (*msr*(*E*) and *mph*(*E*)) were co-localized with aminoglycoside resistance genes (*aph*(*3*”)-*Ib* and *aph*(*6*)-*Id*). Such co-localization suggests mechanisms of co-resistance or the encoding of multidrug resistance, potentially facilitating the presence of co-resistance mechanisms or multidrug resistance capabilities, potentially promoting the survival and emergence of multidrug-resistant *Acinetobacter* strains. The conservation of such ARG clusters indicates that they may be transferred or inherited as a single unit, increasing their persistence and spread within bacterial populations or communities. This phenomenon underscores the potential risk of transmitting multiple ARGs even in the absence of selection pressure on certain resistance determinants.

Furthermore, ARG enrichment was identified within p*dif* modules in plasmids carrying ARGs. The *dif* site comprised a 28 bp DNA sequence featuring two 11 bp Xer protein binding motifs arranged in an inverted repeat configuration, with a 6 bp central region (64). In plasmids, this *dif* site is referred to as the p*dif* module, often occurring in multiple copies throughout the plasmid DNA. Previous studies have reported p*dif* module-associated ARG enrichment in *Acinetobacter* plasmids (64, 65, 66). However, this study, to the best of our knowledge, is the first to document up to 10 different ARGs localized within a single p*dif* module. This observation underscores the coincidence between the presence of ARGs and p*dif* modules in plasmids, emphasizing the critical role of *pdif* recombination sites in the acquisition and accumulation of ARGs in *Acinetobacter* plasmids.

While ARGs are predominantly associated with transposable elements (TEs) and plasmids, their occurrence within prophage regions showed considerable variation across bacterial species. Comprehensive genomic studies of clinical isolates generally report a lack of ARGs in the prophage regions of *Pseudomonas aeruginosa* (67), *Escherichia coli* (68), and *Staphylococcus aureus* (69). However, contrary findings have demonstrated ARG presence within prophage regions of *A. baumannii* isolates (70). These prophage-associated ARGs included beta-lactam (*bla*_OXA-23_, *bla*_NDM-1_, *bla*_ADC-5_, *bla*_OXA-67_, *bla*_OXA-115_, and *bla*_TEM-12_), aminoglycoside (*aac*(*3*)-*I*, *aac*(3)-*Id*, *aacA16*, *aph*(*3’’*)-*Ia*, *aph*(*3*’)-*VI*, *aph*(6)-*Id*, and *aph*(3”)-*Ib*, macrolide (*msr*(*E*) and *mph*(*E*)), and sulfonamide (*sul2*) resistance genes.

In contrast to the findings of Loh et al. (70) and consistent with earlier studies on other bacterial species (67, 68, 69), our analysis detected no ARGs in prophage regions of *Acinetobacter* genomes. This absence may be attributed to the size constraints of phages, which limit their ability to carry accessory genes like ARGs (71). Additionally, the discrepancy with previous studies may stem from advancements in prophage detection tools. Our study utilized the latest iteration of PHASTEST for prophage prediction, which integrates the Prodigal algorithm (72), characterized by reduced false-positive and false-negative rates for open reading frame (ORF) identification. Previous studies employed an earlier version of PHASTER, potentially leading to the detection of ARG-containing prophages that are not confirmed with the updated toolset. This difference highlights the need for methodological standardization in prophage analysis.

In summary, this study highlights the predominant association of ARGs with plasmids in *Acinetobacter* isolates and underscores the significant role of mobilizable and non-mobilizable plasmids, particularly those enriched within p*dif* modules, in driving ARG dissemination. On the other hand, the limited presence of ARGs in prophage regions supports the notion that plasmids are the primary vehicles ARG transfer in *Acinetobacter*. These findings contribute to our understanding of how MGEs mediate antimicrobial resistance and underline the genomic evolution of *Acinetobacter*.

### 4.3. Nested MGE structures as drivers of intracellular and intercellular ARG movement

We analyzed the role of the nested structure of MGEs in the intracellular and intercellular movement of ARGs. ARGs in plasmids were primarily associated with IS elements, transposons, and integrons, consistent with previous findings that ARGs are frequently linked to transposable elements and integrons (17). Although IS-harboring plasmids carrying ARGs accounted for only 8% of the analyzed plasmids, the presence of IS elements highlights their role in facilitating gene shuffling and replication both within and between plasmids.

IS elements can also transfer from plasmids to chromosomes, enabling bacteria to maintain HGT capabilities while reducing the metabolic burden with plasmid maintenance (Horne et al., 2023). IS elements linked to ARGs are key indicators of resistance transfer risk (4, 73). In *Acinetobacter* isolates, IS elements were predominantly found in conjugative MGEs, such as plasmids and ICEs. These elements were enriched in both mobilizable and non-mobilizable plasmids, aligning with our observation of high ARG carriage in these plasmid types. Interestingly, ARG-bearing plasmids had a higher frequency of IS elements compared to plasmids without ARGs, supporting our hypothesis that ARGs are frequently present in plasmids because IS elements, which are highly abundant in plasmids, often harbor ARGs. This aligns with previous studies showing that higher density of IS elements in plasmids is associated with ARG carriage (i.e. carbapenem resistance) (5). Similarly, Che et al. (74) reported that IS elements are more abundant in conjugative plasmids and generally in ARG-bearing plasmids than in plasmids without ARGs. These findings suggest that nested MGEs facilitate hierarchical ARG transfer, promoting the movement of ARGs between plasmids, chromosomes, and even across cells, enhancing overall ARG dissemination.

We also examined the presence of prophages within plasmids, often referred to as “phage-plasmids,” which have been identified in notable pathogens such as *Escherichia, Acinetobacter, Klebsiella,* and *Salmonella* (75, 76). For instance, in *E. coli*, the plasmid p1108-IncY exhibits high nucleotide sequence identity with phage P1, suggesting lysogenization via genetic integration (77). Similarly, prophage regions have been identified in *A. baumannii* plasmids (78). In our study, we detected phage-plasmids in non-*baumannii Acinetobacter* species, such as *A. towneri* and *A. guillouiae*.

Notably, IS elements were abundant in conjugative elements, including prophages within plasmids, particularly in pig waste-associated isolates. IS elements were more common in prophages located in plasmids than those integrated into chromosomes. The nested structure formed by the synergistic integration of IS elements and prophages within plasmids has the potential to further amplify the mobility and transmissibility of ARGs.

### 4.4. Intracellular ARG movement contributes to the formation of putative novel genomic island

Our synteny analysis revealed that the intracellular transfer of multiple ARGs from plasmids to chromosomes plays a significant role in the formation of genomic islands (GIs) variable chromosomal regions that, when enriched with resistance genes, are referred to as genomic resistance islands (GRIs). This transfer is primarily mediated by transposable elements, especially when ARGs are co-localized with transposons or integrons. These findings suggest that ARG movement within bacterial cells (e.g., from plasmids to chromosomes) is a key driver in the formation of new GRIs.

GRIs have been extensively studied in *Acinetobacter*, especially in clinical and pathogenic isolates. In *A. baumannii*, numerous GRIs have been identified and characterized (80–83, 28, 29, 31, 32). However, studies on GRIs in non-*baumannii* species are limited and have mainly focused on clinical isolates from the *Acinetobacter calcoaceticus-baumannii* (Acb) complex, including *A. nosocomialis* and *A. seifertii* (35).

In this study, we identified GRIs in non-*baumannii* strains isolated from pig farm wastewater and discovered novel GRI structures that distinctly differ from those previously reported in clinical isolates (31, 32, 81). Remarkably, this represents the first report of GRIs in non-*baumannii* species outside the Acb complex. The discovery of these GRI structures in non-*baumannii* isolates highlights their adaptation to distinct selective pressures encountered in non-clinical environments, such as those characteristics of pig waste.

Plasmids isolated from pig waste-associated isolates exhibited a notable enrichment of distinct ARG types not commonly observed in patient- and environment-associated isolates. Transposition of these ARGs from plasmid to the chromosome perhaps gives rise to the formation of putative novel genomic resistance islands. These findings highlight unique mechanisms of gene acquisition and genomic organization in non-clinical *Acinetobacter* isolates, expanding our understanding of ARG acquisition and dissemination beyond the clinical context.

## 5. CONCLUSION

This comprehensive analysis of MGEs across *Acinetobacter* species uncovers a complex and dynamic landscape of genetic mobility and ARG distribution. Among the various MGEs, ISs were the most prevalent, followed by plasmids and prophages, with notable differences in their distribution across isolates from patients, environmental samples, and pig waste. Plasmids were identified as the primary carriers of ARGs, with their highest prevalence observed in pig waste-associated isolates. The study identifies several key mechanisms facilitating ARG acquisition and dissemination, including the formation of p*dif* modules and the association of ARGs with transposable elements. The findings suggest that plasmids serve as crucial vehicles for ARG dissemination. While prophages were detected, they did not directly encode ARGs, suggesting a more limited role in ARG dissemination. Overall, this work provides important insights into the mechanisms of ARG transfer and underscores the dominant role of plasmids and their associated MGEs in driving the spread of resistance determinants across diverse environments, enriching our understanding of antimicrobial resistance in *Acinetobacter* species.

## Data Availability

The complete genome and plasmid sequences of all strains reported in this study were deposited in the GenBank database under a BioProject accession number PRJNA1184881.

## ACKNOWLEDGEMENTS

Authors declare no conflict of interest.

This research was performed by the Environment Research and Technology Development Fund (SII-12-1 (3)) of the Environmental Restoration and Conservation Agency provided by Ministry of the Environment of Japan. This research was supported by the Japan Society for the Promotion of Science (JSPS) Grant-in-Aid for Scientific Research (A) (20H00633).

**Kozo Watanabe** is supported by Environment Research and Technology Development Fund (SII-12-1 (3)) and the Japan Society for the Promotion of Science (JSPS) Grant-in-Aid for Scientific Research (A) (20H00633).

**Kenneth A. Bongulto:** Conceptualization, Methodology, Investigation, Formal analysis, Writing-Original draft, Project administration, Visualization. **Ngure Kagia:** Methodology, Formal analysis, Visualization, Writing-Reviewing and Editing. **Hisamichi Tauchi:** Methodology, Writing-Reviewing and Editing. **Satoru Suzuki:** Conceptualization, Methodology, Writing-Reviewing and Editing, Supervision. **Kozo Watanabe:** Conceptualization, Formal analysis, Writing-Reviewing and Editing, Funding Acquisition, Supervision.

## Supplemental Material

Supplementary Figure 1: Frequency of mobile genetic element (MGE) types.

Supplementary Figure 2: Plasmid characterization.

Supplementary Figure 3: Antimicrobial resistance genes (ARGs) within p*dif* modules.

Supplementary Figure 4: Insertion sequence (IS) types in plasmids.

Supplementary Figure 5: Plasmid-associated IS elements.

Supplementary Figure 6: Synteny analysis of plasmids originating from different pig-waste isolates.

Supplementary Figure 7: Synteny analysis of co-residing plasmids from pig-waste isolates.

Supplementary Figure 8: IS element distribution in porphages.

Supplementary Figure 9: Genomic resistance islands (GRIs) in non-*baumannii Acinetobacter* isolates.

Supplementary Table 1: ARG co-localization in plasmids.

Supplementary Table 2: Prophages detected in *Acinetobacter* plasmids.

